# Dermal bone grows by cellular invasion and intercalary biomineralization

**DOI:** 10.64898/2026.09.16.752019

**Authors:** Kate Weymouth-Crocker Jordan, Xintao Zhang, Hiromi Yanagisawa, Marthe J. Howard, David E. Clouthier, Georgy Koentges

**Affiliations:** Laboratory of Genomic Biomedicine and Evolution, School of Life Sciences, University of Warwick,Coventry CV4 7AL,UK; Life Science Center for Survival Dynamics, Tsukuba Advanced Research Alliance (TARA), University of Tsukuba, Ibaraki, Japan; Department of Neuroscience, University of Toledo Health Sciences Center, Toledo, OH 43614, USA; Department of Craniofacial Biology, University of Colorado Anschutz Medical Campus, Aurora, CO 80045, USA

**Author notes:** Correspondence: The authors declare no competing financial interests. Correspondence and requests for material should be addressed to G.K.

## Abstract

Dermal bone architecture bears great evolutionary significance for jawed vertebrates. Its thickness growth is believed to rely on matrix apposition by superficial osteoblasts. To test this hypothesis we employ genetic lineage mapping, conditional gene ablation, intravital matrix labelling and molecular 3D analysis at single cell resolution *in vivo*. We uncover an invasive mechanism incompatible with apposition. The pervasive arrangement of two layers sandwiching a third spongy layer develops from a molecularly defined bi-layer via a previously unknown developmental module: Outer layer osteoblasts form rosette-like assemblies around single invading cells. The latter bear unique molecular signatures and form *de novo* sheets inside the spongy layer, communicating with the layer below. The transcription factor Hand2 organizes this module by orchestrating rosette formation, their molecular heterogeneity, cellular invasion and spongy layer thickness growth. Surprisingly, invading osteoblasts secrete new biomineral matrix inside the older matrix, which keeps expanding. Such intercalary biomineralization provides new perspectives for bone biology and the evolution of endochondral ossification.

## Introduction

The gnathostome craniofacial skeleton is largely composed of dermal bone. Understanding its histogenesis is crucial for elucidating the etiology of many human and animal malformations. Curiously, the cellular and molecular mechanisms of radial/thickness growth in dermal bone are poorly understood^1^. Sutures are places of lateral growth but while dye labelling revealed extensive lateral spread of cells at sutures it remains unknown how cells are incorporated radially into growing bone^2,3^. Sutures close prematurely in human craniosynostosis^4^ and corresponding mouse mutants^5^, but thickness growth continues^6^. This suggests that lateral and thickness growth are genetically separable processes.

The current model for the radial growth of dermal bones is extrapolated from notions of endochondral ossification^4,6,7,8^. It assumes a (cambial) outer layer of osteoblasts depositing new matrix from the outside (appositionally) while osteoclasts^6^ remove bone matrix from the inside, accommodating dermal and brain growth^9^ (see ESM,S1). However, there is no experimental evidence for appositional growth of dermal bone in any vertebrate. In fact, dermal bone formation precedes (bone marrow dependent) osteoclastogenesis both ontogenetically and evolutionarily, necessitating a different mechanism.

Epithelio-mesenchymal interactions are thought to govern dermal bone self-organization^10^. A sandwich of inner and outer compact layers enclosing a spongy (cancellous, vascular) layer have been recognized since Hippocrates^11^ but it has remained enigmatic how this tri-layered organization develops^1^. *Runx2* is known as a dermal bone master-regulator^12,13,1^ expressed in early osteoblasts, while *osteopontin* (OPN) and *osteocalcin* (Ocn) are considered to signify a late osteoblastic cell type, devoid of *Runx2* expression^1,8,14^. Co-expression analysis using RNA in situ hybridization *in vivo* rarely achieves single cell resolution, and the 3D arrangements of dermal osteoblasts have not been studied *in vivo*. It is unclear how well calvarial osteoblasts *in vitro* replicate the spatio-temporal *in vivo* deployment of proteins: *in vitro* these cells do not generate spongy bone. Consequently, bioengineering functional (i.e. trilayered) dermal bone remains challenging.

Here we examine the development of the frontal and clavicle bones, well recognized dermal bones, using immunohistochemistry in three dimensions at single cell resolution in wild-type and *Hand2* mutant mice, and perform *in vivo* birth-dating of nascent bone matrix. We provide the first evidence for intercalary dermal bone growth, affecting the cancellous layer as the major component. We discover rosette-like configurations of osteoblasts as central (and persistent) developmental modules, abolished by cell-specific ablation of the basic HLH transcription factor Hand2 *in vivo*. Rosettes are part of an invasive mechanism by which tightly regulated nuclear translocations of transcription factors and *de novo* cell sheet formation are integrated to accomplish an intercalary biomineralization and remodelling process.

## Results

### From a bilayered to a trilayered dermal bone architecture

Current models of dermal bone formation suggest that dermis vasculature precedes the formation of underlying dermal bone and that an osteoblastic front wraps around pre-existing dermis vessels^15,10^. This notion is not supported by 3D image analysis of CD31+ vWF+ endothelial cells and RUNX2+,OPN+ osteoblasts in the mouse frontal and clavicle at E13-E18. Dermis vessels are not physically connected to the underlying nascent bone, which itself does not harbour any vasculature at its earliest stage of development (Fig. 1a). Instead molecular marker analysis reveals a bi-layered arrangement of Runx2+/Ocn+ cells in outer layer (now termed L1), juxtaposed to OPN+/Hand2+ cells in the inner layer (L3) (Fig. 1b-d). In time these two layers become separated by an intervening space (L2) (Fig. 1b,e-f) that undergoes the most significant expansion of matrix and cellularity among the three. The molecular identity and contiguity of L1 and L3 as bordering sheets is retained throughout development: L1 and L3 encase L2 throughout both a developmental time-course (E13 through P2) and along the naso-parietal axis of frontal bone maturation (Fig. 1g). Marker expression extends from these two layers into the emerging L2 harbouring a new Runx2+/OPN+ cell type (Fig. 1f). Cells carrying the markers for cell (sheet) migration - F-actin, fibronectin (FN) and periostin (POSTN)^16^ penetrate L2 from both sides (Fig. 1h-j,l,n-o) and display hallmarks of the non-canonical *Wnt* pathway usually involved in convergent extension movements^17^ (Fig. 1k).

**Figure 1.**
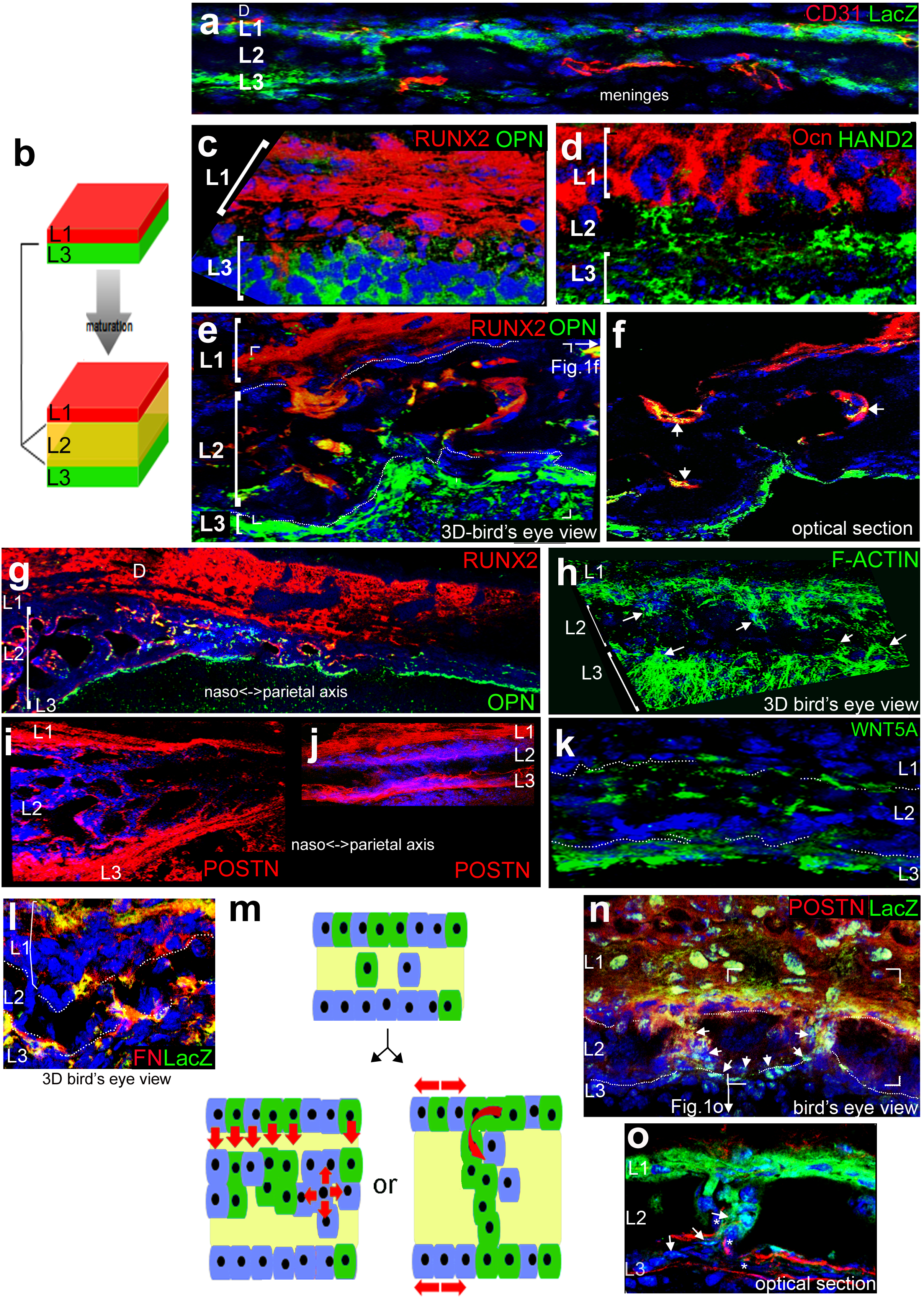
Progression from a bi-layered to a tri-layered molecular dermal bone architecture. Immature frontal bone region (E18) (**a**) is not patterned around dermis (D) vasculature (CD31+endothelia, red; neural crest, green). (**b)** Early dermal bone comprises a molecularly discernible bi-layer at E13 (L1 and L3) that matures into a tri-layered architecture of L1, L2 and L3. (**c, d**) Cells of L1 are RUNX2+/Ocn+/OPN-, cells of L3 are Runx2-/Ocn-/OPN+/HAND2+ as seen by IHC at E13. (**e,f**) The cancellar L2 expands, and is populated by RUNX2+/OPN+ cells (arrows indicates overlap, **f**) at E18. (**e,f,g**) Molecular identities of cells in L1-L3 are retained through a time-course from E13-birth and along the naso-parietal axis of the bone. (**h-j**) Cells in L1 and L3 are highly migratory, as they are immunoreactive (IR) for F-Actin (**h**),Periostin (POSTN) (**i,j**), Fibronectin (FN) (**l**) and invade L2 (arrows in **h**) with little IR at the centre of L2. (**k**) This invasion is further supported by IR for WNT5A, a non-canonical wnt pathway component indicative of a convergent-extension-type process at the L1/2 interface (**l**) The neural crest/mesoderm mosaic (LacZ+/LacZ-) of L1 (**a,l,m,n,o)** in *Wnt1*-cre;R26LacZ transgenic frontals was used to test models of how new cells join L2: (**m**) traditional models (left) imply cells randomly percolating from a generative layer into the matrix^10^, leading to clonal NC or mesodermal cell columns traversing L2 or clonal ‘nests’ within L2 (left); alternatively privileged entry points into L2 would generate strings of NC cells that form new sheets within the matrix (right). 3D reconstruction reveals such strings of LacZ+ NC cells (arrows in **n**) traversing L2 by forming new POSTN+ sheets (see **o** for optical section) connecting L1 and L3. These sheets/clasps are of mixed composition but dominated by NC (*, LacZ-). Original magnifications: **a-f, h,j-o** 63-100x; **g, i** 10x. Nuclear DAPI, blue.

The frontal bone was previously fate-mapped to be of pure NC origin^18^. Surprisingly, more detailed genetic analysis (*Wnt1*-Cre;ROSA26R mice) reveals that the nascent bone L1 is a mosaic of neural crest and mesoderm, while early L3 is only mesodermal (Fig. 1a,l,n,o). This afforded the opportunity to test the model according to which osteoblasts randomly percolate in sheet-like fashion from the cambial layer into the underlying nascent (osteoid) bone matrix^9,10^. If that were to be the case we would expect radial columns of NC versus mesodermal cells traversing L1-2 or L3-2 boundaries or clonal nests of β-galactosidase (β-GAL)+ NC cells within an otherwise mesodermal L2 (Fig. 1m). We do not find evidence for these. Instead, at earliest stages sheets of migratory (POSTN+) NC cells start from specific points within L1 and extend across L2 into L3 so that the initially thin and sparse L2 (Fig.1a,d) becomes increasingly populated (Fig. 1n-o). (See **ESM_S1-S3** for model comparison and marker choices).

### A new histogenetic module: osteoblastic rosettes and clasps

To investigate the morphogenesis of the L1-L2 connection points, we employed 3D reconstructions at single cell resolution. At the apices of these connection points we discovered rosette-like arrangements of 5-6 RUNX2+ cells (Fig. 2). Within a given rosette one cell traverses the L1/L2 boundary and displays nuclear RUNX2 (Fig. 2a-b,e,f-g,h). In the most immature condition, this single nuclear-RUNX2+ cell extends a cellular process and contacts a cell at the L2/L3 boundary (Fig. 2b, arrow), a configuration we term a clasp. This nuclear-RUNX2+ cell is also characterized by nuclear localization of HAND2, a transcription factor known to bind RUNX2 under *in vitro* conditions^19^, while HAND2 remains cytoplasmic in the remainder of the rosette (Fig. 1e). The nuclear-RUNX2+/HAND2+ cell is also OPN+ and Col1A1/A2+ (Fig. 2f-h). This molecular heterogeneity of rosettes is retained even in the most mature bones. The connection between rosettes and cellular sheets within L2 is never broken. As only a single cell provides the physical connection to the underlying L2 clasp/sheet, our 3D analysis shows that L2 sheets form *de novo* and not by invaginating L1 sheets. This suggests that rosettes and clasps are key components of a hitherto unrecognized generative architecture governing the growth in thickness of dermal bones.

**Figure 2.**
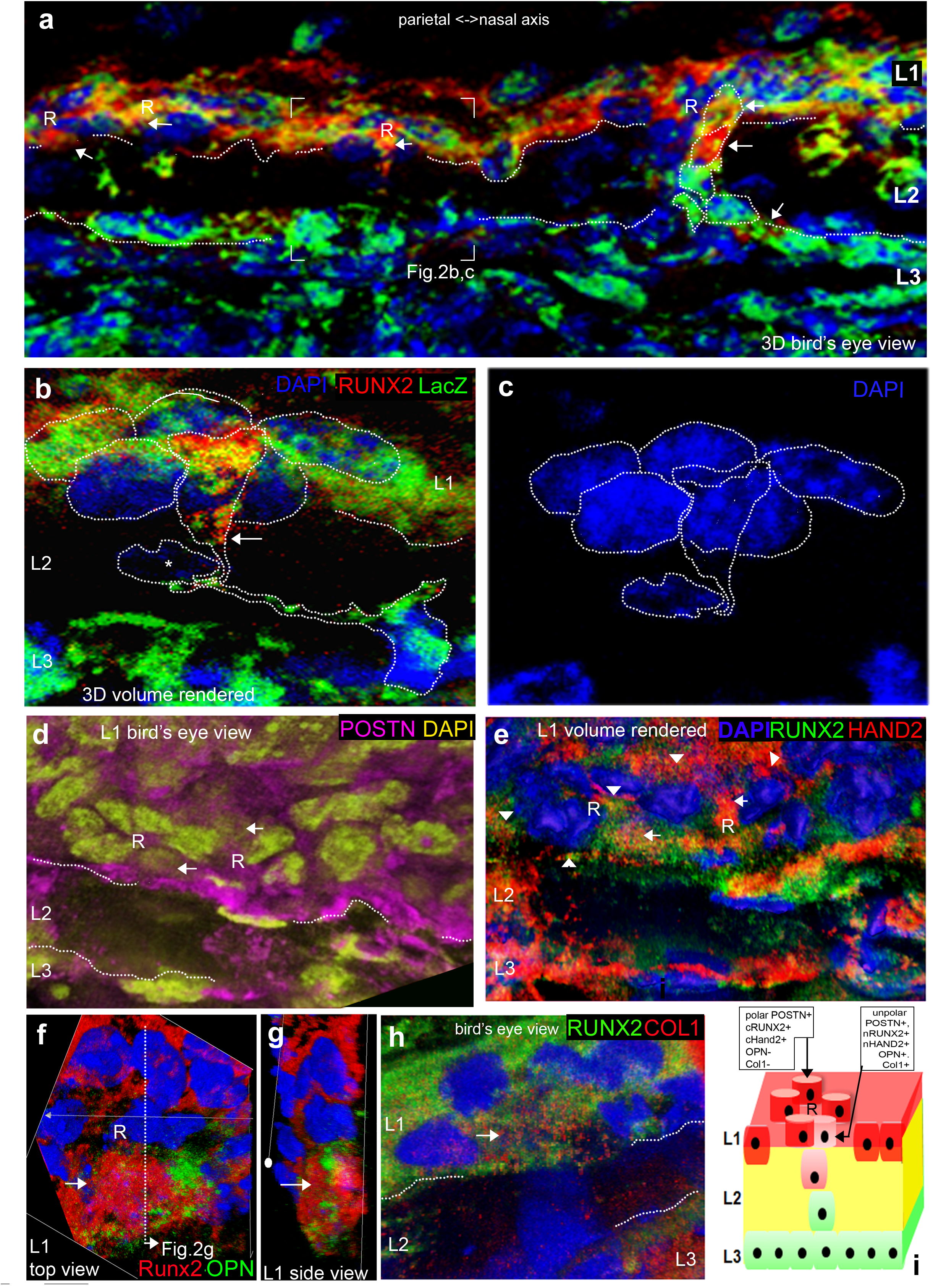
Osteoblastic rosette architectures in dermal bone and its molecular heterogeneity. L1 to L2 entry points are organized as cellular rosettes. (**a**) 3D surface views along the naso-parietal maturation axis of the frontal bone (left to right) reveal a rosette (R) of 5-6 cells at the apex of each clasp (**b-h**); clasps interconnecting L1 and L3 comprise either single cells (**b,c**) or increasing numbers of cells (right, arrows in **a**). The majority of cells in a rosette display cytoplasmic RUNX2 (**b, e-f, g-h**), POSTN (**d**), and HAND2 (**e**), cytoplasmically polarised toward L2 (arrowheads in **e**). A single cell in each rosette displays nuclear RUNX2 (arrow, **a-h**), nuclear HAND2 (**e**), OPN (**f** and corresponding nuclear cross-section **g**), and Collagen I (**h**) and reaches into L2 (**b,c)**. (**i**) The nRUNX2/nHAND2/OPN/Col1+ signature is also carried by cells within L2 clasps (arrow, **a**; pink cell **i**) but not by mesodermal cells (* in **b**). Rosettes are present from E13 (**f-g**) through birth (**d**) and in both the frontal bone (**a-c,f-h**) and clavicle (**d-e**). All 63-100x. Nuclear DAPI in blue (**a-c, e-h**) or gold (**d**).

### Hand2 controls rosettes and L2 architecture

The remarkable co-localization of nuclear (n) RUNX2 and nHAND2 within the single invading rosette cell prompted us to test whether NC-specific Hand2 ablation affects jointly rosette architecture and L2 elaboration *in vivo*. Contrasting with the highly cancellous wild-type condition, *Hand2fl/fl;Wnt1-Cre/R26R* (which we refer hereafter as *HAND2*^*cko*20,21^) embryos have no L2 growth: the thin plane of mineralized matrix lacks any trabeculation in both the frontal and clavicle bones (Fig. 3a-b). We can identify cell-autonomous effects of neural crest-specific *Hand2* ablation in the L2 mosaic as the *Hand2* mutant cells are also β-GAL+ (Fig.3d,h).

**Figure 3.**
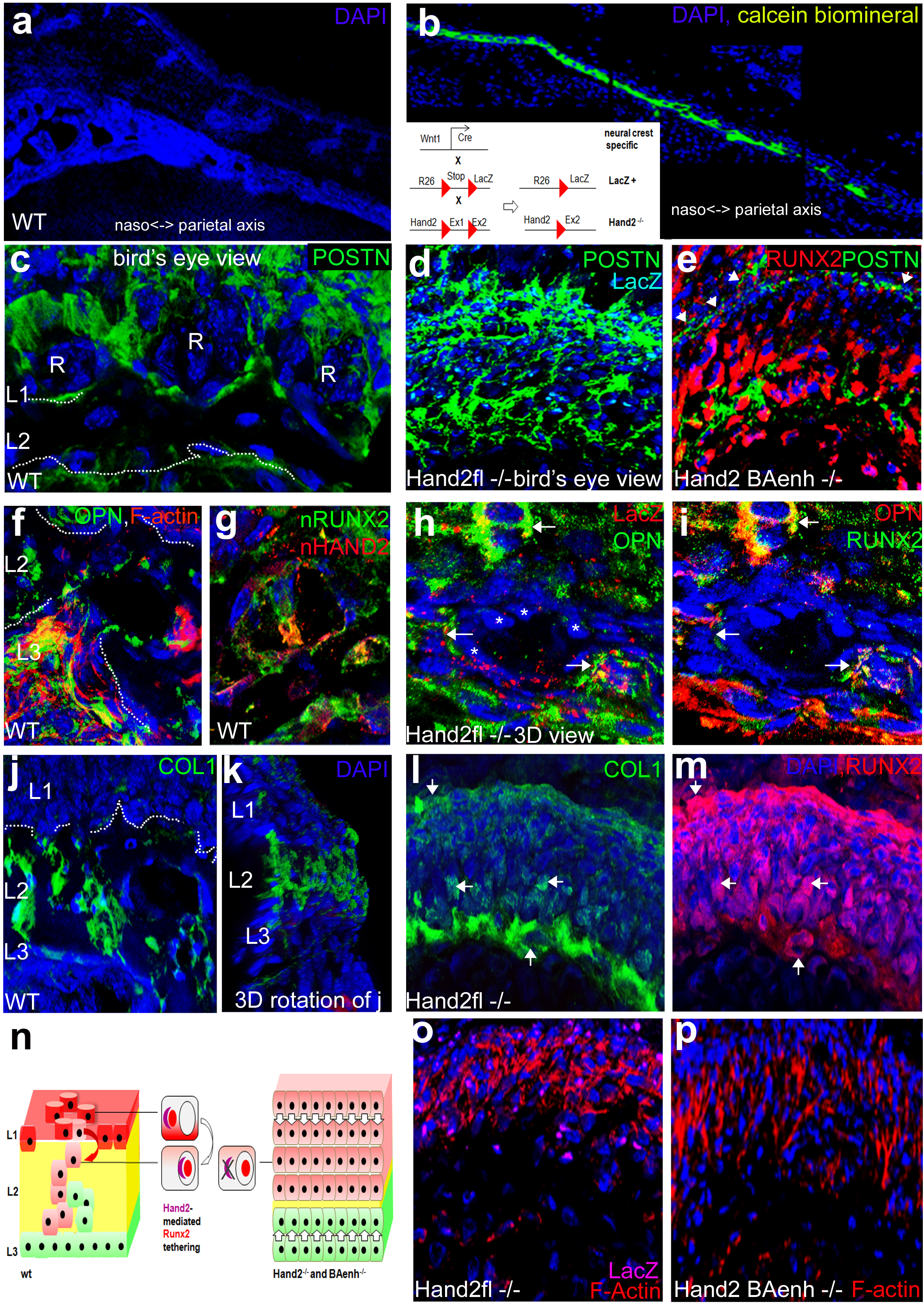
Hand2 mutants lack rosettes and an cancellar L2. In contrast to the highly trabeculated wild-type frontal bone at E18 (**a**), NC-specific deletion of HAND2 (**b**, inset) results in a frontal bone with a single thin Calcein+ mineral sheet along the naso-parietal axis. (**c**) Wild-type clavicle rosettes (R) are nested in depressions made of cytoplasmically polarized POSTN, visible in 3D surface view. (**d**) Rosettes are absent in Hand2-/-mice where POSTN polarity is lost in LacZ+ (blue) cells that now randomly traverse an unorganised L1-L3 space. (**e**) Hand2BAenh mutants phenocopy this behaviour: RUNX2-nuclear cells randomly fill the area bounded by a POSTN+ seam (arrowheads, **e**). In contrast to flat, migratory OPN+ (**f**), nRUNX2/nHAND2 (**g**) wild-type cells, Hand2 mutant (LacZ+) OPN/RUNX2+ cells (arrows in **h,i**) remain blast-like and do not join sheets of flattened mesodermal (*) cells. (**j,k**) Collagen I is confined to L3,L2 clasps- and single nRUNX2/OPN+ rosette cells in wild-type while in the mutant (**l**) the clasp-less hybrid L1/L2 layer is completely filled by Col1+/nRUNX2+ cells (see arrows, **l-m**). (**n**) This suggests that HAND2 tightly and conjointly regulates RUNX2 nuclear localization, rosette and clasp formation in wild-type, a behaviour mediated by the branchial arch enhancer for dermal bone formation. (**o-p**) Hand2 mutation (by conditional Hand2 removal or branchial arch enhancer ablation) results in a dense, columnar L1/2 organisation akin to traditional dermal bone models (Fig.1m,left), as indicated by ubiquitous radial F-Actin fibres. Stage-matched frontal (E18) **a,b,f,h-i**; clavicle (newborn) **c-e,g,j-p. a-b** 10x; **c-p** 63-100x. Nuclear DAPI, blue.

In a bird’s eye (surface) view, L1 displays polarized POSTN localisation demarcating wild-type rosettes (Fig. 3c). These are no longer visible in *Hand2*^*cko*^ mutants (Fig. 3d,e). Instead, all cells within L1 (as indicated by a retained POSTN seam, Fig. 3e arrowhead) display nuclear RUNX2, abolishing the molecular heterogeneity distinctive of wild-type rosettes (Fig. 3e). The architectural distinction between L1 and L2 is lost in mutants where β-Gal+/POSTN+ cells indiscriminately traverse L2 with no discernible clasps (Fig. 3d). OPN+/RUNX2+/HAND2+ (β-GAL+) NC cells are flattened and form sheets within a wild-type L2 (Fig.3f,g). However, they remain blast-like and individualized and do not join into nascent clasps formed by mesodermal (β-GAL-) cells (Fig. 3h-i, arrows) in *Hand2* NC mutants. In wild-type bone, strong Col1A1/A2 reactivity is initially confined to L3 and the single RUNX2/OPN+ cell of the rosette (Fig. 2h,3j); as more cells participate in clasps the Col1A1/A2 pattern extends to follows the clasp (Fig. 3k) This does not take place in mutants, where strong immunoreactivity remains confined to the external margin of L1 and the L2/3 interface (Fig. 3l, arrows). Col1A1/A2 and POSTN thus border the hybrid L1/L2, which is full of migratory (F-actin+) nuclear-RUNX2+/OPN+ cells (Fig.3m,o-p) but is devoid of rosettes and clasps.

The ubiquitous nuclear localisation of RUNX2 in *Hand2*^*cko*^ (β-GAL+) NC cells points to an unusual cytoplasmic tethering mechanism by which HAND2 regulates RUNX2 within rosette cells (Fig. 3n). In cells with cytoplasmic HAND2 we find RUNX2 trapped in the cytoplasm. When HAND2 is nuclear, RUNX2 is nuclear as well. In Hand2 mutant cells RUNX2 translocates into the nucleus independently. Thus, Hand2’s cytoplasmic localization overrules thenatural tendency of RUNX2 to translocate into the nucleus. The same mutant phenotype is found when HAND2 is ablated through targeted excision of its branchial arch (BA) enhancer (Fig. 3e,p)^22^. This provides genetic evidence that rosettes and L2 clasps are dependent upon BA-enhancer driven *Hand2* function, without which the (NC derived) dermal bone regions lack L2 growth.

### Intercalary biomineralization of dermal bone

In order to relate the newly discovered rosette/clasp architecture to the mode of L2 biomineralization, we performed a (E14-E18) time course of Calcein stains for biomineral. This reveals discrete biomineralization stages (Fig. 4a-c): 3D analysis shows that the earliest Calcein+ areas within L2 are islands (yellow in Fig.4g) separated from one another by rosettes (R) and sheets of cells (grey in Fig.4g) communicating between L1 and L3 (Fig. 4a,d,g). With time these islands coalesce laterally and are intersected horizontally to form parallel sheets (green in Fig. 4b,e), followed by radial mineral interconnections between the sheets (Fig. 4c). At the latest stage, the outer mineral surfaces are separated from L1 and L3 by unmineralized clasps (Fig. 4f-g). These significant biomineral shape changes require a remodelling that is not commensurate with an appositional model of successive matrix deposition from the outer side. They rather suggest cellular invasion/sheet formation and subsequent biomineral resculpting. We tested this hypothesis *in vivo* by intraperitoneal injection of two fluorochromes (Calcein and Xylenol Orange (XO)) into time-pregnant dams at different time points of bone development (Fig. 4h). The dyes incorporate into newly deposited matrix and permit us to birthdate matrix spatially. As controls we measured dye retention times in the maternal and embryonic system and performed dye swaps to control for possible differences in dye diffusion/uptake kinetics. We carefully calibrated the assayed *in vivo* emission spectra by confocal microscopy to obtain non-overlapping fluorescent signals (see supplement for details). As unincorporated dyes are cleared 2 days post injection, we injected Calcein at E14 and XO at E16. We reconstructed in 3D bone regions with varying degree of L2 biomineralization at E18 or P2.

**Figure 4.**
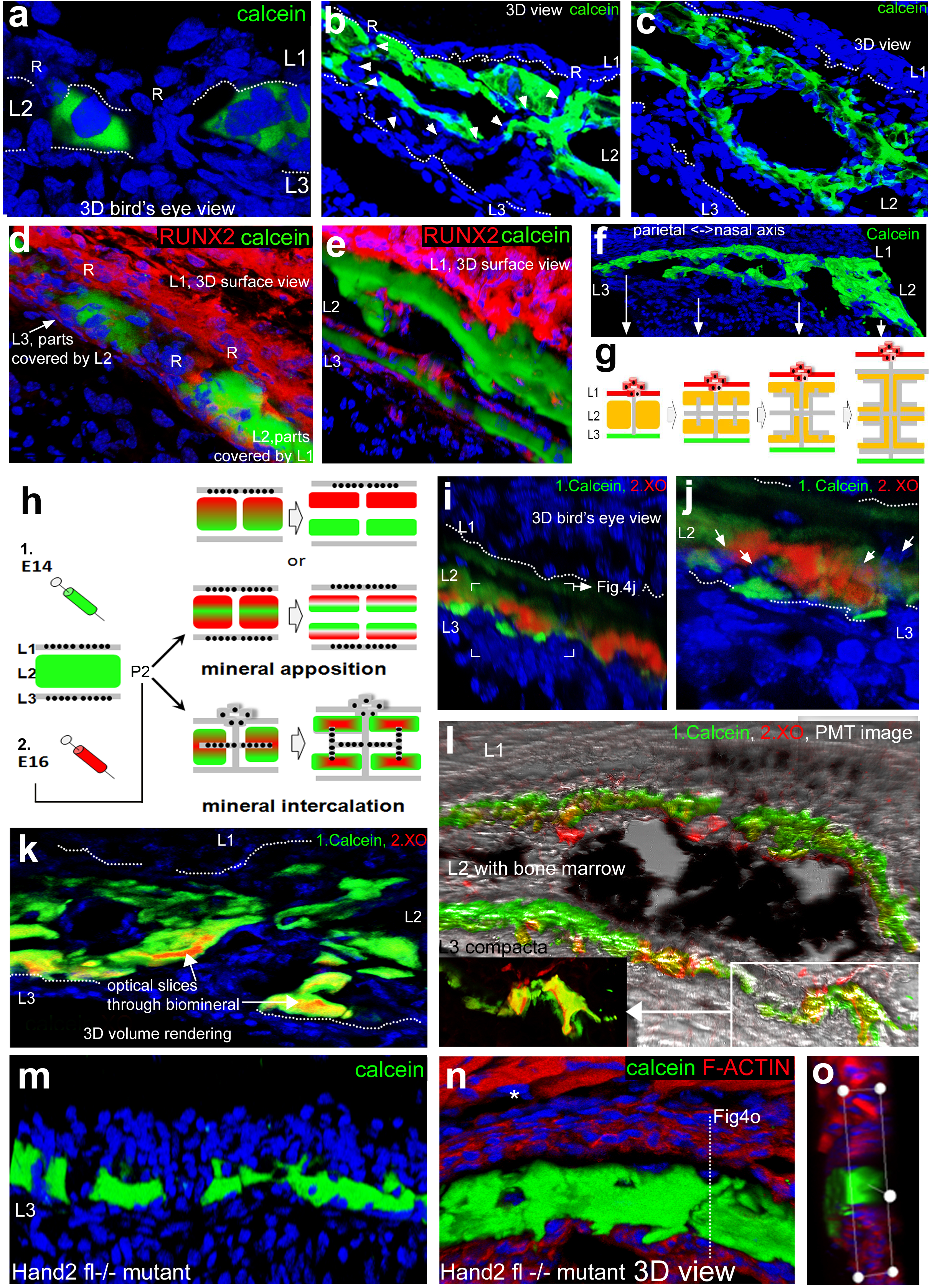
Intercalary growth of L2 biomineral *in vivo*. During a developmental time-course the mineral of the frontal bone progresses from islands, to parallel horizontal planes which are vertically connected at a late stage (histological calcein addition from E14-E18, **a-c**). Mineralized areas are separated by sheets of cells in L2 (arrowheads **b**), connected to L3 and rosettes in L1 (R, rosette in bird’s eye view of L1 in **a,d**; **e**). (**f,g**) This process proceeds along the naso-parietal axis and implies extensive remodelling of mineralized areas. (**h**) *In vivo* matrix double labeling experiments test appositional versus intercalary growth mode of L2. (**i-l**) Optically sectioned 3D reconstructions show that the second label (XO) is always nested inside mineral labelled by the first dye (calcein), incompatible with appositional growth (sections **i-j**; 3D reconstructions **k-l**). Thus, cells must have invaded (green) L2 before the second label was incorporated. This labelling pattern persists inside the most mature regions of the bone at P2 (**k-l**), where no evidence for (osteoclast-mediated) matrix catabolism from the bottom can be found. Comparison between 4 i and 4l shows massive expansion of initially labelled green layer, incompatible with apposition. Labelled matrix corresponds to genuine mineral (seen by PMT imaging, **l**). Interstitial biomineralization is not achieved in Hand2-/-mutants who lack cancellar mineral deposition and L2 vasculature in the frontal (**m**) and dermal NC-derived clavicle (**n** and corresponding cross section **o**; * NC-derived muscle attachment region, see **ESM,S3** for details. **a-e, i-k, m-o** 63-100x; **f, l** 10x. Nuclear DAPI, blue.

In an appositional model the matrix deposition by osteoblasts/osteocytes on one or both sides of L2 would lead to the second colour (XO, red) topping or sandwiching the first (Calcein, green) (Fig. 4h). Contrasting this, a model of intercalary growth, where cells invade into previously laid down matrix before depositing new matrix, would result in the second colour being nested inside the first (Fig. 4h). Indeed, a bird’s eye view of immature L2 regions shows the red label fully nested within green sheets of earlier matrix (Fig. 4i-j). In more cancellated areas, the newer (red) matrix is always confined to the inside of the older (green) matrix while the overall thickness of L2 has increased significantly (Fig. 4k). We observe no topping or sandwiching of the earlier (green) by the later (red) matrix. The incorporated dye-coloured matrix persists into the fully matured condition displaying well known mineral features (Fig. 4l). This is remarkable because it necessitates the earliest laid down matrix to expand very significantly with time. We see no erosion of early matrix label from the inner side of the bone, as current models of osteoclast-mediated erosion would predict.

As rosettes and clasps within L2 are abolished in *Hand*^*cko*^ embryos, we examined frontal and clavicular biomineralization in these mutants. *Hand*^*cko*^ embryos exhibit only a narrow sliver of mineral without cancellar growth, similar to the most immature wild-type mineral (Fig. 4m-o, similar to Fig. 4a). This suggests an arrest at the earliest stage of L2 biomineralization. New matrix-secreting osteoblasts are unable to form L2 sheets that we propose to act as radial growth and biomineralization scaffolds.

## Discussion

We have uncovered a novel generative process and developmental module governing thickness growth in mammalian dermal bones (ESM,S1). 3D time course analysis by combinatorial IHC *in vivo* allowed us to trace the transition from a two-layered to a three-layered architecture (comprising L1,2,3). Molecular marker signatures (Osteocalcin+/Osteopontin+) widely associated with a late osteoblastic cell type^1,8,19,23^ are found at the earliest stages of dermal bone development, which challenges their validity as late differentiation markers(ESM,S3). The molecular signatures of L1 and L3 persist, indicating that protein deployment is linked to cellular position inside the growing bone rather than to differentiation (ESM,S2).

L2 is dominated by a previously unrecognized nuclear-RUNX2/OPN+ cell type. This cell type emanates from rosettes within L1 that display a distinctive and stereotypic molecular heterogeneity at single cell resolution. Osteoblastic rosettes have not been seen previously, they are only discernible in 3D reconstructions *in vivo*.

We do not know if the single nuclear-RUNX2/OPN+/Coll1+ cell within a rosette results from a dynamic singling-out process or acts as a permanent guidepost for other cells. Resolving this would require intravital *in utero* imaging, currently impossible. 3D analysis also shows that cellular invasion does not happen by L1 invagination but by a single cell ‘bee-lining’ at rosettes and *de novo* sheet formation within L2. How could such bee-lining be accomplished? In many developmental contexts convergence-extension (CE)-type movements and polarization of cell shape are accomplished by the non-canonical Wnt/ PCP pathway^17,24^. Interestingly, a central component of this pathway, Wnt5a is also active here (Fig. 1k). The inability of *Hand2* mutant cells to flatten up and partake in sheet formation after invasion places Hand2 genetically into this pathway and points to a growing number of CE/Wnt/PCP genes evolutionarily co-opted into vertebrate skeletogenesis^25,26^.

Many layers of Runx2 post-transcriptional control have been described^8,14,27^. We show cytoplasmic RUNX2 tethering by HAND2 within rosettes as a new *in vivo* phenomenon, invisible to traditional RNA ISH analysis: In *Hand*^*cko*^ mutants, the cytoplasmic RUNX2 retention, rosettes, cell polarization, L2 cancellar elaboration and biomineralization are conjointly abolished (Fig.3). This identifies *Hand2* as the first gene to orchestrate this newly discovered generative architecture. It will be interesting to determine the genetic mechanisms organizing these developmental modules within the L1 plane of the growing bone as this will sculpt its outside relief. The cellular invasion observed (Fig.1l-o) also predicted an interstitial biomineral growth mechanism to explain the extensive remodelling we see during development (Fig.4g). *In vivo* double pulse labelling of dermal bone matrix, which has not been done to date, proves that new matrix is deposited on the inside of older matrix, incompatible with traditional apposition. The accompanying paper describes this L2 growth mechanism in molecular detail. The generative architecture documented here provides a new framework for re-analyzing many mutants and for understanding a host of craniofacial pathologies^6^ (see ESM,S1). A detailed molecular comparison will elucidate whether and how the phylogenetically older dermal ossification programmes^28,29^ were evolutionarily co-opted for endochondral ossification processes whose analysis has so far dominated skeletal developmental biology.

## METHODS SUMMARY

Multiplex (single, double, triple and quadruple) fluorescent immunohistochemistry was performed on mouse specimen from E13 – P2 using standard protocols. Image acquisition was performed using Leica TCS SP2 and SP5 systems at 10x-100x, as z-series with an average step size ∼0.6µm. 2D image analysis was conducted using LeicaLiteAS and ImageJ; all 3D z-series reconstructions (surface and volume rendering; optical slicing) were generated using BioimageXD (http://www.bioimagexd.net). Neural crest lineage analysis was done using a *Wnt1*-cre transgenic strain crossed with a Cre-reporter R26R-LacZ^30,31,32^. Analysis of the effect of Hand2 ablation in neural crest cells was undertaken using a Hand2 conditional knockout in which a floxed exon1 is excised when Hand2fl/fl mice are crossed with *Wnt1-*cre;R26LacZ mice ^20,21^; the simultaneous presence of the conditional mutant Hand2 and reporter transgenes allowed us to monitor cell-autonomous effects in these mosaic tissues (made of neural crest and mesoderm/rhombomere 1-neural crest). Phenotypes were compared to that of mice in which the Hand2 branchial arch (BA) enhancer is deleted via homologous recombination, the BA*enh*-/-mice^22^. A biomineralization time-course was established using *ex vivo* histological addition of Ca^2+^-chelators calcein or xylenol orange at 5mg/mL in conjunction with immunohistochemistry and PMT imaging. *In vivo* labelling was conducted under HOL PPL 70/7178. Intraperitoneal injections of each agent (calcein 10mg/kg, xylenol orange 90 mg/kg) into time-pregnant C57BL/6J dams were done at E11-18 (with a minimum two day gap between injections). Specimens were isolated for analysis at E18 - P2(see full methods for details of all antibodies and protocols in **ESM_S4**).

## Supporting information

Jordan1 SupplementLast

## Supplementary Information

ESM,S1-S4 is attached.

## Acknowledgements

We thank Andrew Lumsden, Nick Dale and Jonathan Millar for critical reading of the manuscript, Dan White for technical assistance with BioImageXD, Samantha Dixon and Ian Bagley for expert animal husbandry and care, B. Ryll for the histological preparation of 3 slides, Ian Portman for imaging support. The work was funded by ConquerChiari, Human Frontiers and the Wellcome Trust (all awarded to GK), NIH (R01HD0648240)(H.Y). NIH grants DE018899 (to DEC) and NS040644 and DK067064 (to M.J.H.).

## Author contributions

KWJ and GK designed all experiments and wrote the manuscript. KWJ performed and analyzed all experiments. DEC, MJH, HY and X.Z. generated and provided essential transgenics. All authors contributed towards data analysis and the final manuscript.

## Author information

The authors declare no competing financial interests. Correspondence and requests for material should be addressed to G.K.

