## Supplementary material for "Dermal bone grows by cellular invasion and intercalary biomineralization": Jordan1 SupplementLast

### **SUPPLEMENTARY TEXT**

**S1: Comparison of traditional appositional and our model**

**S2: Cellular identity is positionally, not temporally defined**

**S3: Molecular marker choice**

**S4: Detailed methods/technical appendix**

### S1. CONTRASTING THE TRADITIONAL AND INTERCALARY MODELS OF DERMAL BONE THICKNESS GROWTH

#### Traditional model

Traditionally the lateral growth of dermal bones is believed to be a direct result of sutural growth whereby (pre-)osteoblastic progenitor cells within sutures migrate laterally and contribute the cellular material that will later deposit matrix in an apposition fashion<sup>1,2</sup>. FGFR signalling has been involved in the timing of sutural growth and closure and defects/mutations in this pathway have been implicated as causative for craniosynostosis and related phenomena<sup>3-5</sup>. Interestingly, in these conditions and in corresponding mouse mutants it was shown that sutures close prematurely while thickness growth continues<sup>6</sup>. In patients this leads to clover-leaf shaped compensatory deformations of the skull, replicated in mouse mutants. As suture patency does not directly affect thickness growth this indicates separate cellular and molecular mechanisms might regulate lateral and thickness growth of dermal bone. The exact mechanisms of thickness growth are currently unknown with apposition being the assumed (only) one.

The assumed mechanism of thickness growth in dermal bone has largely been inferred from what is known about endochondral ossification, which has been subject of the most rigorous molecular analysis over the past two decades<sup>7-9</sup>.

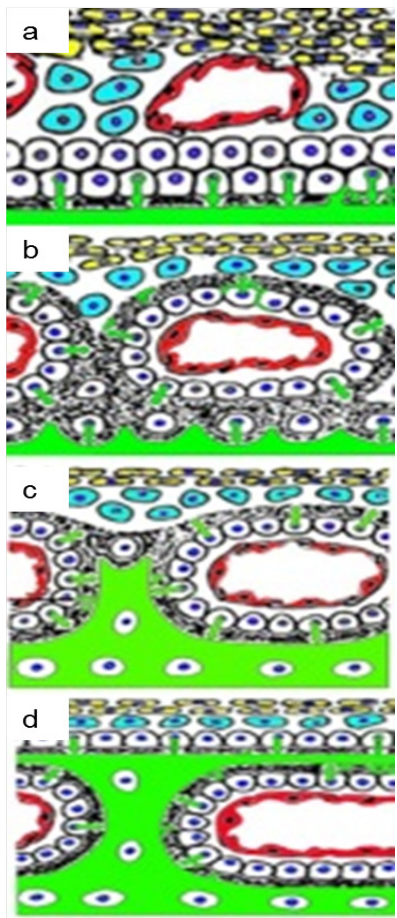

In the traditional model of thickness growth during intramembranous ossification, dermal bone formation is preceded by an increase in vascularisation at the site of future skeletogenesis, after which mesenchymal cells migrate to the area and are subjected to epithelial-mesenchymal interactions<sup>10</sup>. These interactions lead to the condensation of the mesenchyme into trabeculae<sup>11</sup>. Proper osteogenesis initiates when a sheet of mesenchymal cells in the trabecle differentiate into the osteoblastic lineage, as indicated by the expression of genes associated with a pre-osteoblastic condition, namely Runx2 and Collagen I<sup>12</sup>. Next, a single layer of osteoblastic precursors further differentiate from pre-osteoblasts into osteoblasts (a transition reportedly indicated by the expression of Osteopontin, and BSP II) and these cells start producing osteoid (Fig. S1a), which mineralises to form the hardened calcified tissue<sup>7,12</sup>. Some osteoblasts percolate at random from the generative front and become engulfed by the matrix they secrete and further differentiate into osteocytes capable of small amounts of rapid osteolysis<sup>13</sup> (Fig. S1b); at this stage the next layer of pre-osteoblasts differentiates into osteoblasts and a second layer of osteoid is deposited (Fig. S1b). This implies that a front of deposited (green) matrix needs to become connected with other matrix that has been secreted by osteoblasts surrounding blood vessels in more superficial layers. Thus one expects to find (albeit transient) discontinuities of matrix at the future connection points between old and new (appositionally deposited) matrix.<sup>8</sup> (Fig. S1c-d). This means that pre-existing blood vessels become engulfed by newly deposited bone matrix in time.

Simultaneously, it is assumed that bone marrow-derived osteoclasts are introduced to the system via the vasculature. These are capable of eroding the bone from the bottom to accommodate the lateral growth of the underlying brain while new matrix is deposited at the top.

In this model, at a given time-point there is only a single generative layer of fully differentiated osteoblasts (which abuts the pre-osteoblastic layer). This cambial arrangement would result in the earliest deposited matrix being found in innermost position, while the superficial most matrix being the most recently deposited one. If, during bone formation, different dyes that incorporate into freshly deposited matrix are injected one would expect a layered distribution with the first dye colour to be found at the base and the later injected dye to be incorporated more superficially.

Any further remodelling to the bone (to accommodate additional vasculature, bone resorption, etc.) is accomplished via osteoclasts arising from the bone marrow from monocyte/megakaryocyte precursors<sup>14</sup>. Unlike endochondral bone formation, dermal bone develops from the mesenchyme without any cartilage precursor as a template.

The architecture of the intramembranous anlagen is, therefore, dictated by two features in the traditional model: the pre-existing vasculature, around which the bone must form, and the single outer generative layer of osteoblasts secreting the osteoid and mineralising the matrix in an appositional fashion (Fig. S1). The growth of the inner/vascularised/cancellar layer that has been observed since Hippocrates, has so far not been studied mechanistically.

2400 years ago Hippocrates wrote:

Oeuvres Completes D'Hippocrate. Hippocrates. A. Littre. Amsterdam. Adolf M. Hakkert.

The bone at the middle of the head is double, the hardest and most compact part being the upper portion, where it is connected with the skin, and the lowest, where it is connected with the meninx (dura mater); and from the uppermost and lowermost parts the bone gradually becomes softer and less compact, till you come to the *diploe*. The diploe is the most porous, the softest, and most cavernous part. But the whole bone of the head, with the, [p. 146] exception of a small portion of the uppermost and lowermost portions of it, is like a sponge; and the bone has in it many juicy substances, like caruncles; and if one will rub them with the fingers, some blood will issue from them. There are also in the bone certain very slender and hollow vessels full of blood. So it is with regard to hardness, softness, and porosity.

Translation: Hippocrates. De capitis vulneribus/Peri ton kephale tromaton.

Part1. Translation: The Genuine Works of Hippocrates. Hippocrates. Charles Darwin Adams. New York. Dover. 1868.

http://www.perseus.tufts.edu/hopper/text?doc=Perseus%3Atext%3A1999.01.0250%3Atext%3Dvc%3Asection%3D1

However it was assumed to arise through angiogenic sprouting from external and internal vasculature<sup>15,16</sup>.

### Intercalary growth model:

In contrast to the traditional model of dermal bone development, described above, current results have supported an alternative intercalary growth model of dermal bone thickness growth. The principal features of the new model are described in detail below.

#### Two versus 1 generative layers

In contrast to the expectations dictated by the traditional model, findings presented here propose an alternative interstitial model of dermal bone formation favouring intercalation of new cellular material from two generative surfaces as the mode of thickness growth. In the intercalary model the external envelope, comprising molecularly distinct L1 and L3, is established by E13 (Fig. S2a). These two layers contribute cells to the embellishment and radial growth of the intervening space, L2 which grows in thickness as shown by our mosaic analysis of neural crest fate mapping experiments. L2 houses the mineralised matrix and cancellous core of the bone (Fig. S2).

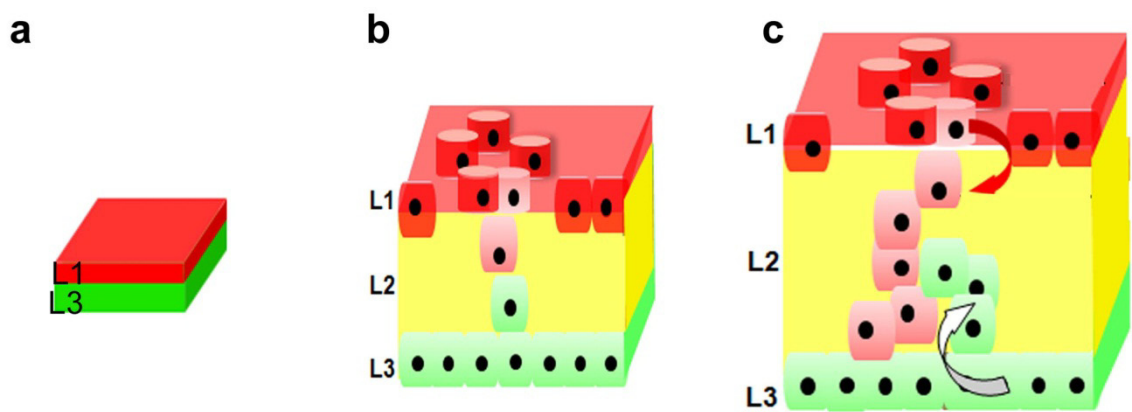

#### Layer 1,3 contiguity and thickness growth of L2

The establishment of the two generative layers in the frontal bone (L1 and L3) by embryonic day E13 indicates a molecular heterogeneity of the morphologically homogenous mesenchymal condensations and an early pre-patterning of the frontal bone primordium (Fig. S2a). The expression pattern and morphology of the two generative layers continues through bone development and birth (Fig. S2a-c). The integrity of the molecularly defined Layers 1 and 3 is not broken at any stage of development, essentially creating a closed system. Because of this observed L1 and L3 contiguity we found that the elaboration of the cancellar L2 constitutes the most significant component of overall anatomical thickness growth.

### **Blood vessels form as part of L2 elaboration and do not precede it**

By using 3D analysis in developmental time courses we were unable to find evidence for L1 and L3 wrapping around pre-existing dermal vasculature as posited by the traditional model. If involution or wrapping of these layers around pre-existing dermal blood vessels did occur one would expect to find at least grooves of partially engulfed dermal vasculature, which were never observed.

### **Rosette-like entry points of cells**

In the traditional model of apposition cells from the generative (cambial) layer percolate individually and randomly into the underlying matrix. Our genetic mosaic analysis shows that cells do not display such behaviour. Instead the entry points of cells in L2 were highly constrained to areas where cells in L1 are organised into rosette structures. Such rosette structures are only visible once 3D reconstructions have been performed and are not visible in traditional analysis of histological sections (Fig. S2b-c).

Rosettes are found in single embryonic time slices and across ontogeny. L2 grows in thickness as additional cellular material is added to it from generative layers L1 and L3. In the most immature stage of frontal bone development where a tri-layered architecture (L1-L2-L3) is already evident (by E16), L2 is a cell sparse region that does not contain any blood vessels. As maturation progresses L2 is increasingly populated. 3D reconstructions reveal sheets of connected cells dictate the cancellar internal organization of L2 (Fig S3, below).

### **Intercalary versus appositional matrix deposition**

The continuous intercalation of new cellular material into L2 has extensive ramifications for the deposition of biomineral in dermal bone (Fig. S3).

The traditional model of appositional growth would have predicted that newly deposited matrix is found superficial to older matrix, a hitherto untested assumption. Our double dye labelling reveals the opposite. The biomineralized areas are expanded from cells intercalating and increasing the mineralised area by expanding the earlier matrix scaffold from the inside. Mineralization first takes place at E14 (Fig. S3) as soon as the tri-layered architecture is established. Distinct islands are separated by unmineralized sheets of cells. These islands coalesce to form parallel sheets that continue to be separated by cells maintaining contact with the generative layers (Fig S3). The differing structure of biomineralized areas between immature and mature regions of a dermal bone implies a remodelling process to take place (the accompanying paper details the molecular processes involved in this).

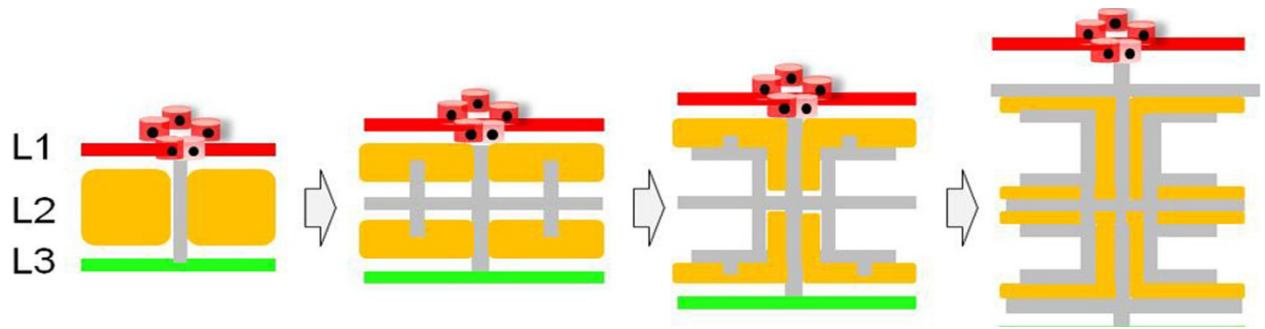

### **S2. CELLULAR IDENTITY IS POSITIONALLY NOT TEMPORALLY DEFINED IN DERMAL BONE**

A key tenet of the traditional model is that the maturational sequence from pre-osteoblasts to osteoblasts to osteocytes would have to be reflected in the spatial location of these cells within the bone. This means that the most immature cells would be localized most superficially and a maturation gradient should be reflected by expected marker distributions. .

If that were the case we would expect the following (as detailed in the figure S4 below, based on expression patterns expected from previously published studies):

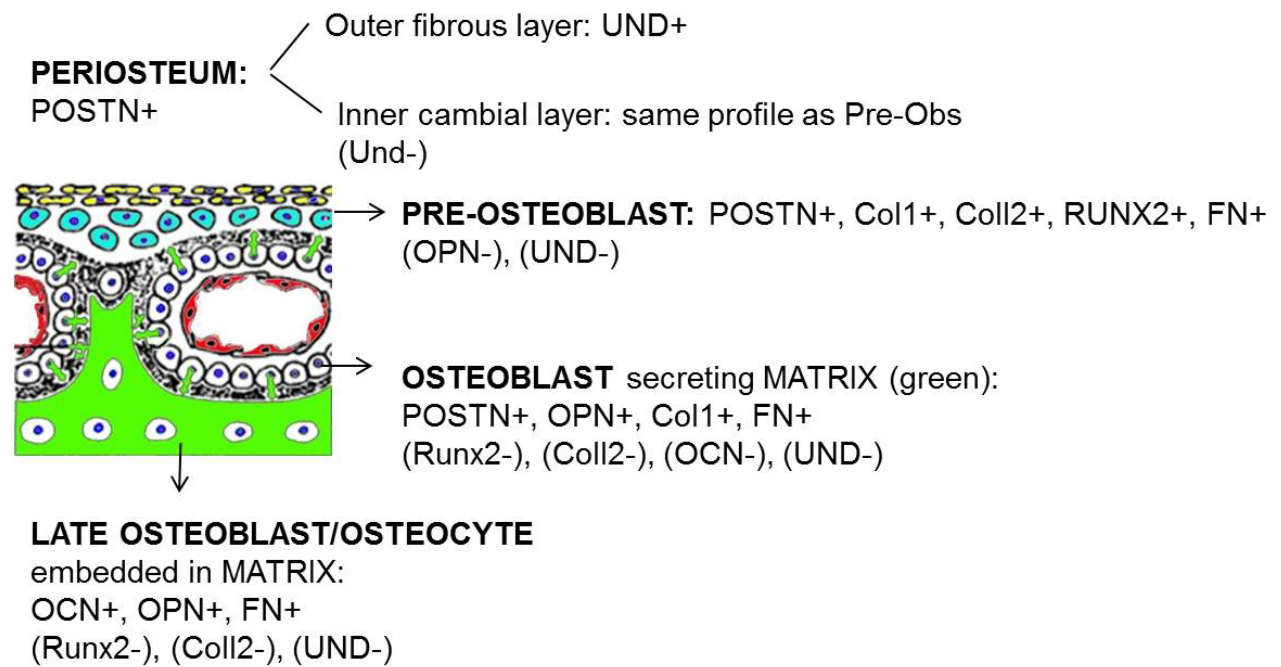

1. Periosteal markers such as Periostin (POSTN) would only be found in the superficial Layer 1. We see POSTN encasing L2 from both sides (in L1 and L3).

2. Undulin is expected in the most superficial fibroblastic layer of the periosteum<sup>12</sup>. We find Undulin immune-reactivity in all three layers including positivity within osteoblastic cells of L2.

3. Preosteoblasts would be expected to express POSTN, Collagen IA2 (Coll1), Collagen II (Coll2), Fibronectin (FN) and Runx2 but not markers of mature osteoblasts, such as Osteopontin (OPN) or Osteocalcin (OCN)<sup>12,17,18</sup>. We observe that cells of Layers 1 and 3 have unique marker expression retained across a time-course analysis. Cells of L1 are Runx2/Coll2/OCN+, while cells of Layer 3 are OPN/Coll1+ and cells of both layers are immune-reactive for POSTN and FN.

4. Loss of Runx2 and Col2, gain of OPN and no overlap between Runx2 and OPN was expected. Using *in vivo* 3D IHC analysis of these proteins we detect a significant fraction of Runx2/OPN double positive cells within L2. This was unexpected as Abzhanov, on the basis of performing *in situ* hybridization of dissociated dermal bone cells in the chick did not detect such significant overlap<sup>12</sup>. We speculate that this population was missed because it would be very difficult to efficiently dissociate these cells from their surrounding matrix. Furthermore single cell resolution *in vivo* is required to precisely map the nature of the overlap.

5. Late osteoblasts and osteocytes are supposed to be POSTN-/Col1- but positive for OPN and OCN. We already observe OCN/Runx2+ cells within the dermal layer and within rosettes. This suggests that Osteocalcin cannot be utilized as a *bona fide* marker for late stages of osteoblast differentiation as is commonly assumed in experiments of calvarial cell cultures.

These experimental findings, done at single cell resolution, are not compatible with traditional expectations of cambial growth, a simple maturational gradient and its accompanying appositional matrix maturation patterns.

#### S3. CHOICE OF MOLECULAR MARKERS

The current study was aimed to be an inclusive examination of protein deployment during dermal bone thickness growth *in vivo*. A wide-array of proteins were analysed using IHC such that we could simultaneously probe bone maturation and resolve widely conflicting expression domains of these markers as reported in dermal bone literature<sup>9,12,17,19,20</sup>.

Most importantly, only when we were able to identify a given marker protein on the inside of a given cell (using 3D reconstruction and optical sectioning) would we determine it to be expressed by that cell. This is an essential step to verify the molecular identity and cell population diversity at single cell resolution in wildtype and mutant animals.

Based on the **traditional model of dermal bone development** (see above), we chose to analyse the following markers:

The presence of the vasculature, assumed to pre-pattern the bone, was examined with endothelial markers CD31 and von Willebrand factor (vWF) and pericyte marker α-SMA<sup>21-23</sup>. The formation of the mesenchymal anlagen would be monitored using Fibronectin (FN), a cell adhesion molecule and marker of sheet migration downstream of TGFβ1 required for mesenchymal condensation formation, secreted by cells depositing a basal membrane<sup>24</sup> and expressed by early pre-osteoblasts<sup>17</sup>.

Traditional pre-osteoblastic markers also included Runx2 (also widely expressed in fibroblasts), as well as Collagen I and Collagen II (expression analysis in **accompanying paper 2**)<sup>17</sup>. The previously identified transition from pre-osteoblasts to mature osteoblasts capable of matrix secretion/mineralization would be marked by Osteopontin gain and Runx2 loss (**Abzhanov, Komori** review). The final maturation of osteoblasts and differentiation into osteocytes would traditionally be monitored by mapping Osteocalcin expression<sup>17</sup>.

Presence of the periosteum would be indicated by immunoreactivity of Undulin (Collagen XIV)<sup>20</sup> and Periostin (POSTN); notably, POSTN is reportedly expressed by pre-osteoblasts and has been found to regulate sheet migration of cells, a phenomenon well described for neural crest cells migrating into branchial arches or into the developing heart outflow tracts<sup>17,25,26</sup>. Most importantly, POSTN is a member of the FasciclinII family of proteins involved in coordinating cellular behaviours via homophilic interactions, ranging from *Drosophila* to vertebrates (**ref** doi:10.1016/j.modgep.2005.03.005, Chiba et al. Nature 374, 1995, 166-168, ).

Given Runx2's critical role in bone development<sup>27-30</sup> we also chose to analyse a newly defined regulator of Runx2, Hand2<sup>31</sup>; we were especially interested in the abnormal skeletal phenotype previously observed, but uncharacterized, in strains of Hand2 mutant mice (details of the mutant strains, as well as additional images of the mutant phenotype are presented in Figure S5, below) as their detailed histology and layer morphogenesis has not been studied in any species.

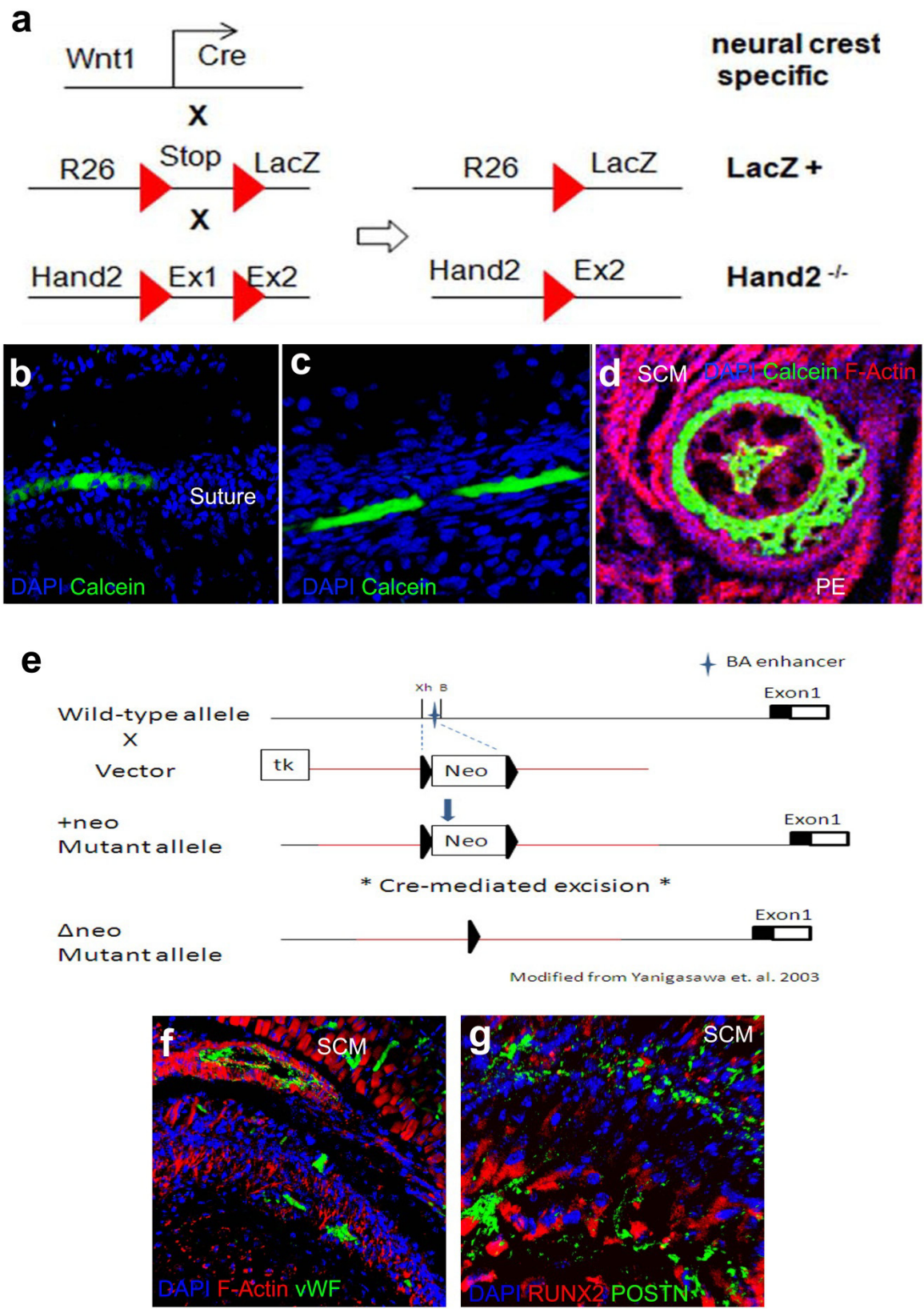

Furthermore, given the expected lateral migration of cells at the margins of the forming bone and the apparent sheet-like behaviours of cells spanning L2, we undertook the additional examination of F-Actin and components of the non-canonical *Wnt* signalling pathway, including Wnt5a.

Mixed neural crest and mesodermal origin of frontal bone.

Finally, previous fate mapping studies have generated conflicting reports as to the lineage origins of the frontal bone<sup>32,33</sup>. The location of the neural crest:mesoderm boundary within the neurocranium has been heatedly disputed, with the margin of the neural crest territory alternatively hypothesized to be the frontal:parietal interface<sup>34</sup> or the parietal:occipital junction<sup>32</sup>. The composition of the frontal bone has been argued to be entirely neural crest<sup>32</sup> or primarily mesodermal<sup>33</sup>. Notably, Couly *et al.*, who argue for an entirely neural crest-derived frontal bone, employ a standard grafting technique (quail-chick chimera) in their analysis, however, the final time-point of their study (just over mid-way through incubation) preceded definitive osteogenesis in the frontal and parietal areas examined by several days. Moreover, no molecular markers were used to identify the cells as osteoblasts and , conversely, mesodermal derivatives were not labelled separately either. As such, their fate maps were generated at stage similar to E13 in murine development, as presented above, where the mature architecture of the frontal bone is not yet established and any late invasion or contribution of mesodermal cells could not be analysed. Conversely, Evans and Noden maintain there is a large contribution of the mesoderm to calvarial tissues and their analyses occur at a slightly later time-point than Couly; however, their use of targeted injection of replication-incomplete retroviruses is not a *de rigueur* technique due to potential concerns over cross mesodermal:neural crest contamination.

Establishing a time-course of the contributions of the neural crest and mesoderm to both the generative layers (L1 and L3) and the fully formed L2 is a key component of understanding the cellular origins and histogenesis of the frontal bone, a bone for which homologies across vertebrates are well established<sup>35</sup>. As such, in addition to mapping the generative architecture of dermal bone formation, the current study also aimed to resolve the widely contested issue of lineage origins of the frontal bone using genetic lineage analysis (see Methods for details).

### S4. Detailed Methods

#### Generation of transgenic animals

***Wnt1*-Cre x ROSA26LacZ:** Neural crest cells were permanently labelled utilising the well-characterised *Wnt1* dorsal neural tube enhancer region. The *Wnt1* enhancer confers transgene expression on the entire dorsal neural tube neural crest precursor population, and their subsequent progeny (with the exception of NC-cells from rhombomere I; McMahon & Koentges, unpublished)<sup>1</sup>. *Wnt1*-Cre mice were crossed with Cre-reporter mice carrying transgenic floxed resistance-pA cassettes in front of LacZ under the control of ROSA26<sup>2</sup>; this combination yields offspring in which the floxed cassettes are subject to Cre-mediated excision. The neural crest cells of these *Wnt1*-Cre x R26-LacZ offspring can be visualised with either X-gal staining (LacZ)<sup>3</sup>, or labelled with an anti-βGal/LacZ antibody. *Wnt1*-Cre mice were obtained from the Jackson Laboratory (USA) and crossed with the ROSA26 reporter mice also obtained from Jackson Laboratory (USA).

**Conditional knockout, *Hand2*<sup>fl/fl</sup> x *Wnt1*-Cre-R26LacZ,** bred and maintained by H. Yanigasawa at the University of Texas (Dallas): *Hand2* was specifically deleted in neural crest cells using the following method: briefly, loxP sites were inserted into the *Hand2* allele flanking exon 1 in the 5' UTR<sup>4</sup>. *Hand2*<sup>fl/fl</sup> mice display a normal phenotype and have normal *Hand2* expression, validated by qRT-PCR<sup>5</sup>. Excision of *Hand2* and visualisation of the cells in which the excision has taken place, is achieved by crossing the *Hand2*<sup>fl/fl</sup> mice with hemizygous *Wnt1*-Cre and homozygous ROSA26LacZ mice.

**Targeted deletion of *Hand2* BA enhancer,** bred and maintained by D. Clouthier at the University of Colorado: *Hand2* expression in the branchial arches requires the Endothelin-1 (ET-1) enhancer. This 208 bp enhancer was deleted using homologous recombination<sup>6</sup>. The enhancer lies within a 754 bp *XHOI*-*BamHI* fragment, which was replaced with a loxP-flanked *neo* cassette. Cre-mediated excision results in *BAenh*<sup>-/-</sup> mice.

#### Immunohistochemistry

Samples were prepared as above, with the following modifications: specimen from embryonic stage E16 onwards were fixed overnight in fresh 4% PFA at 4°C. All specimen for immunohistochemistry from embryonic stage E16 onwards were cut to a thickness of 12-17 µm onto charged slides, from E10-E14 specimen were cut to a thickness of 10 µm. Multiplex immunohistochemistry was performed on sections of embryos from all crosses listed above. Slides were defrosted at room temperature for 5 min, fixed in 4% paraformaldehyde for 15 min at room temperature, rinsed 3 x in PBS and blocked with permabilisation for 1 hour at room temperature. As a default, 20% Roche Western Block Solution (WBS) (Roche) + 0.1% Triton in PBS was used as the default blocking solution. Following blocking, primary antibodies were incubated at 4°C overnight in the blocking solution. The next morning samples were washed for 1-2 hours(s) with the wash solution (10% blocking solution without any detergent), with changes every 5-15 min. Secondary and conjugated antibodies were then incubated for 1 hr at room temperature in the Blocking Solution, followed by 1-2 hour(s) of washes (with frequent changes) and a PBS rinse. Slides with tyramide amplification (β-Galactosidase (AbCam) and Hand2 (R&D) antibodies) had the following additional steps: quenching of endogenous peroxidases with 0.3% H<sub>2</sub>O<sub>2</sub> during blocking; tyramide amplification (Perkin Emler Kit) with 5 μL conjugated (FITC, Cy3 or Cy5) tyramide in 500 μL Ampli-Buffer for 10-15 min; stopping of amplification reaction with 4% PFA or 0.01 N HCl for 10 min and 3 x PBS rinse. All slides were counterstained with DAPI (Invitrogen), administered either at 1:1000 for 5 min, or 1:2000 for 20 min with Rhodamine Phalloidin (Chemicon 1:250). Slides were then rinsed a final time in PBS and mounted in Mouviol + DABCO. Slides were cover-slipped and sealed with nail varnish. Details of primary antibodies used can be found below. All secondary antibodies were obtained from Invitrogen and used at a concentration of 1:200.

All primary antibodies are commercially available and have been extensively validated for specificity and cross-reactivity by the respective commercial sources and the community, using immunohistochemistry (with and without antigenic peptide pre-treatment), Western blotting and various other methods. We have also performed controls of secondary antibody non-specific binding by omitting the primary antibody. Double positivity for respective molecular markers is only scored and accepted if we have seen **intracellular** immuno-colocalization using 3D volume rendering at single cell resolution.

### PRIMARY ANTIBODIES

| Protein | Host | Working Dilution | Company | Reference |
| --- | --- | --- | --- | --- |
| β-Catenin | Mouse, IgG1k | 1:100 | Millipore | 05-665 |
| β-Catenin | Mouse, IgG1 | 1:100 | BD | 610153 |
| β-Galactosidase | Chicken, IgY | 1:500+Amp<br>1:1500 | AbCam | ab9361 |
| CD31 (PECAM) | Rabbit, IgG | 1:20 | AbCam | ab28364 |
| Collagen I | Mouse, IgG1 | 1:200 | AbCam | ab6308 |
| Collagen I | Rabbit, IgG | 1:100 | AbCam | ab21286 |
| Collagen I | Rabbit, IgG | 1:100 | AbCam | ab59435 |
| Col14A1 (Undulin) | Mouse, IgG1k | 1:250 | Stratech | H0000737-M01 |
| Fibronectin | Rabbit, IgG | 1:100 | Sigma | F3648 |
| Hand2 | Goat, polyclonal | 1:50+Amp1:1500 | R&D | AF3876 |
| Notch (activated) | Rabbit, IgG | 1:100 | AbCam | ab8925 |
| Osteocalcin | Mouse, IgG1 | 1:100 | Stratech | NB600-1528 |
| Osteopontin | Rabbit, IgG | 1:100 | AbCam | ab63856 |
| Periostin (POSTN) | Rabbit, IgG | 1:100 | AbCam | ab14041 |
| Runx2 | Mouse, IgG2a | 1:200 | AbCam | ab76956 |
| Wnt5a | Rabbit, IgG | 1:100 | AbCam | ab72581 |

#### ***Ex vivo* matrix labelling**

Matrix Labelling: histological addition of matrix mineral labelling agents in conjunction with immunohistochemistry was conducted prior to slide mounting following the immunological staining or in its absence. Mineral matrix of bone was stained with a labelling agent, either: Calcein, Xylenol Orange, Doxycycline, Calcein Blue, or Alazarin Complexone (All Sigma). Labelling agents were made into solution with PBS, with 5 mg/50mL an effective concentration of all agents to allow visualisation of labelling (for spectral acquisition see below). Slides were dipped in the labelling reagent for 10 sec – 45 sec (agent dependent), rinsed several times with PBS then mounted as described above.

#### ***In vivo* matrix labelling experiments**

Animals obtained by continued use were used to map a time-course of bone mineralisation. Labelling agents were chosen that could be visualised without any treatment to enhance signal in emission areas with non-overlapping spectra such that they could be concomitantly analysed (for controls and details of image acquisition, see below). Concentrations of labelling agents were chosen such that the labelling was expected to be conferred to the dam and her offspring, without any pharmacological side effects<sup>7</sup>, as determined in a host of previous studies on long bones<sup>8-10</sup>: calcein 10 mg/kg and xylenol orange 90 mg/kg. Labelling agents were administered via intraperitoneal injection to mouse dams. Dosages were sufficient to confer the agents to the fetuses to achieve embryonic labelling. Mineralised bone was permanently labelled at 2 time-points between E11 and P0. Dosages were established from literature values on studies of bone-repair in long bones as above, and were prepared in Phosphate Buffered Saline (PBS, 137 mM NaCl, 2.7 mM KCl, 8 mM Na<sub>2</sub>HPO<sub>4</sub>, 1.4 mM KH<sub>2</sub>PO<sub>4</sub>, pH 7.4) and sterilised prior to administration. All protocols were conducted in accordance with Home Office Licence PPL70/7118 (Analysis of the kinetics of dermal bone mineralization). Specimen subjected to the protocol were terminated via an appropriate Schedule 1 method and the tissues were harvested (no sooner than 3 days after the last injection). Samples were fixed in fresh 4% PFA overnight, embedded in OCT (Optimal Cutting Temperature) media (TissueTek, VWR) and stored at -80°C until cryosectioning. Cryosectioning was performed on OTF 5000 Cryostat (Bright, UK) at a thickness of 15-17 µM. Samples were counterstained with 1:1000 DAPI (Invitrogen) and Image Acquisition and Analysis were conducted as per details below.

**Time-course of dye retention (FigM1).** To determine for how long the dye is available for matrix incorporation after injection into pregnant dams and to confirm the individual labelling agents would clear the system prior to the addition of the next compound we performed the following control experiments. A single labelling agent was added to a dam between E11-12 and the offspring were isolated at two time-points, E18 and P2. Matrix mineralisation first occurs at E14, thus any labelling agent from the E11-12 injection that was retained in the system for more than 2 days would incorporate into the matrix.. Analysis revealed the matrix of the animals injected between E11-12 did not contain any labelling agents, when examined at E18 and P2. This control confirms the dye does not circulate within the system for more than 2 days and incorporation times closely matched the injection times. Thus, a 2 day gap between injections was chosen as the desired minimum injection period between addition of different dyes in the later experimental phase. Examining the two time-points of sacrifice was done to ensure that any areas that were labelled by the E11-12 injections were not missed due to subsequent remodelling *in vivo* (leading to the erroneous conclusion that the dye had not been retained through the mineralisation).

**Dye swap.** To ensure that different fluorescent dyes would not label different types of matrix preferentially and would not differ significantly in their ability to penetrate tissues, dye swap experiments were performed: Each labelling experiment was conducted as a pair involving a dye reversal; ie calcein injection at E14 and xylenol orange at E18 has the dye swap control of xylenol orange injection at E14 and calcein at E18. Patterns produced by the labelling regime and the dye swap were compared and were found to be the same.

**Image Acquisition:** Confocal microscopy was performed using Leica TCS SP2 and SP5 systems using 10x, 20x, 40x, 63x, and 100x lenses. Alexa Fluor 488 was excited at 488 nm and emission was measured around 500 nm on the FITC channel; Alexa Fluors 546, 555, RPh and Cy dye 3 were excited at 543 nm and emission measured around 555 nm (TRITC channel). Alexa Fluor 647 and Cy dye 5 were excited at 633 nm and emission measured around 655 nm. DAPI stain was excited at 364 nm and emission measured around 400 nm. In order to relate molecular markers with the directionality of collagen fibre systems we performed reflectance imaging by exciting with the 488 laser and collecting using PMT. PMT imaging has traditionally been used in the field to detect the directionality of collagen fibre systems due to the birefringence of that molecule.

The detection range of emissions was optimised to guarantee no overlap or bleed-through between the channels (FigM1A). All image acquisition occurred in the x, y and z-planes, resulting in a z-series. The average z-series comprised a step size of ~0.4-0.9 $\mu\text{m}$ and the entire depth of the section was imaged. Brightfield images were acquired under a dissecting microscope, in a single optical plane.

#### Image Acquisition: Matrix Labelling (FigM1)

Samples that had been subjected to mineralised matrix labelling were imaged and the emission spectra were empirically determined by confocal microscopy. This allowed us to optimise the detection parameters for all matrix dyes, ensuring that no spectral overlap occurred.

| Labelling Agent | Excitation/Emission | SP5 Laser | SP2 Laser | Overlaps: |
| --- | --- | --- | --- | --- |
| Alizarin complexone | 530-580 / 624-645 | DPSS 561 | HeNe 1.5 mW (543) |  |
| Calcein | 495 / 517 | Ar 100 mW (488) | Ar 100 mW (488) | FITC |
| Calcein blue | 373 / 420-440: | Diode 20 mW (DAPI) | Diode 20 mW (DAPI) | DAPI |
| Doxycycline | 390-425 / 520-560 | Diode 20 mW (DAPI) | Diode 20 mW (DAPI) | RPh |
| Xylenol orange | 440/570 / 610: 555 | DPSS 561 | HeNe 1.5 mW (543) |  |

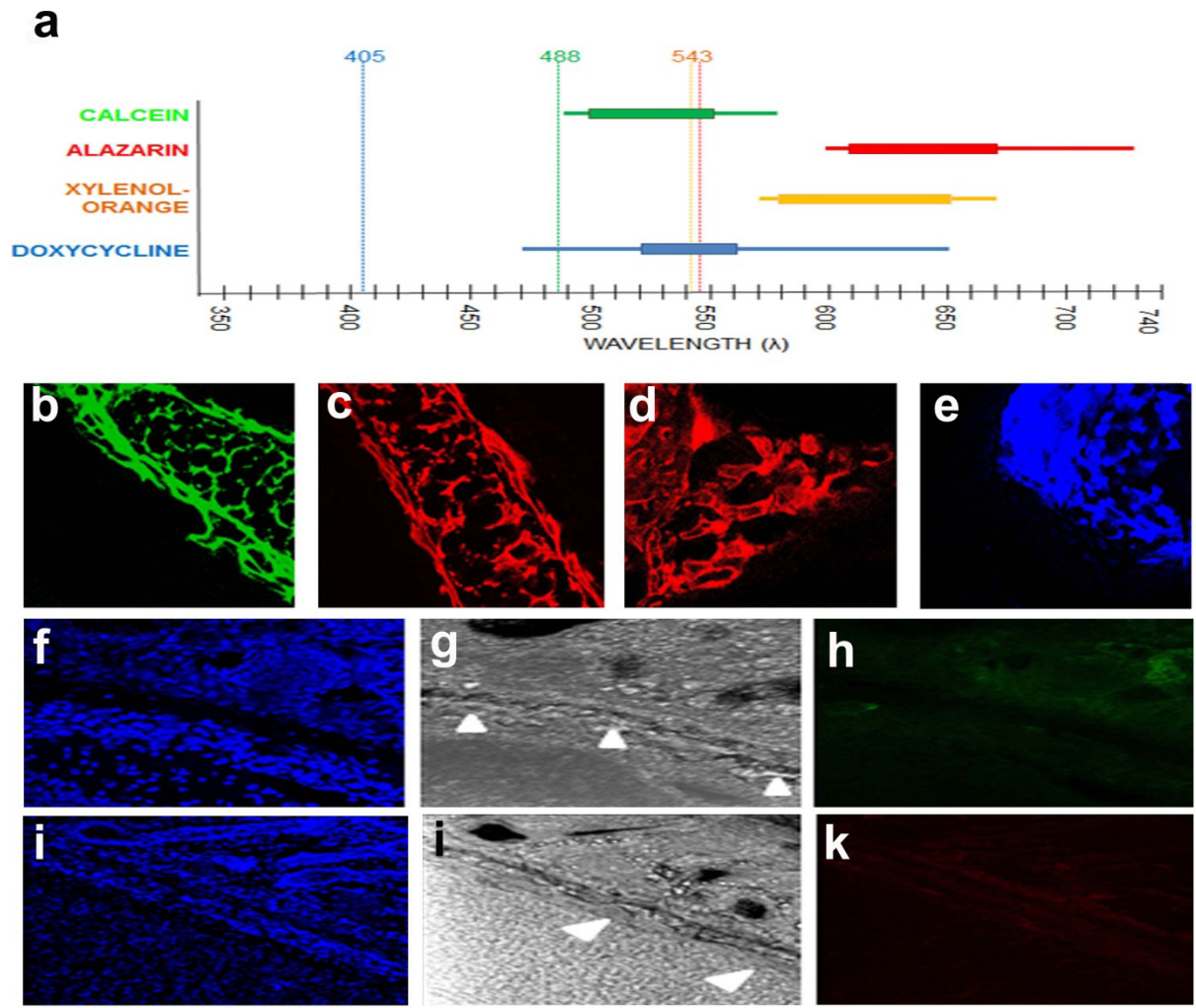

**Figure M1: Controls for *in vivo* mineral labelling experiments.** (a) Empirically determined emission spectra for confocal microscopy: the recommended *in vivo* dosage was histologically added to sections of wild-type, newborn mice. Maximal detection range (wavelength) range indicated by line, OPT (optimal range for detection) indicated by bar. Emission spectra is significantly wider than reported in the literature. (b-e) Images collected in regions of endochondral ossification from OPT range for calcein (b), alizarin (c), xylenol orange (XO) (d), and doxycycline (e); calcein and XO were used in experiments due to their good imaging quality, non-overlapping OPT range, and specificity for mineralised areas. (f-k) Sagittal sections of a wild-type frontal bone at postnatal day 3 (P2), following injection of a single labelling agent at embryonic day E11 after conception. Images are oriented cranial to the top and ventral to the right. Nuclear counterstain DAPI (blue). No labelling agents (calcein, h; XO, k) are detected in the bone, therefore the labels cleared the system before the matrix mineralised at E14. Reflection image (g,i) show mineralised areas along the frontal bone (arrowheads). In subsequent experiments, areas of labelling from compounds injected at E14 were visible at P2, confirming the absence of labelling in the control is due to dye clearing not remodelling of the matrix.

### **Image Analysis**

**Image Analysis, 2D:** 2D image reconstructions were completed using LeicaLiteAS, ImageJ and Adobe Photoshop software. Images were either of individual optical sections from a z-series, or collapsed maximal or average projections of z-series. Image files from the confocal were converted into colour and channels were superimposed.

**Image Analysis, 3D:** All 3D reconstructions (surface and volume rendering) of z-series were created using the freeware BioImageXD developed by the Universities of Jyväskylä and Turku in Finland and the Max Planck Institute of Germany: World Wide Web http://www.bioimagexd.net. We thank Dan White for unwavering support and help with running this programme.
